# Cholesterol biosynthetic pathway induces cellular senescence through ERRα

**DOI:** 10.1101/2021.10.20.465100

**Authors:** Dorian V. Ziegler, Mathieu Vernier, Joanna Czarnecka-Herok, Charlotte Scholtes, Christelle Machon, Jérôme Guitton, Jennifer Rieusset, Vincent Giguère, Nadine Martin, David Bernard

**Affiliations:** Centre de Recherche en Cancérologie de Lyon, Inserm U1052, CNRS UMR 5286, Centre Léon Bérard, Université de Lyon, Lyon, France; Goodman Cancer Research Centre, McGill University, Quebec, Montreal, Canada; Biochemistry and Pharmacology-Toxicology Laboratory, Lyon-Sud Hospital, Hospices Civils de Lyon, F-69495 Pierre Bénite, France; CarMeN Laboratory, INSERM U1060, INRA U1397, Lyon, France; Departments of Biochemistry, Medicine and Oncology, McGill University, Montreal, Quebec, Montreal, Canada

**Author notes:** Equal contributions (Nadine Martin, David Bernard).

**Keywords:** Cellular senescence, ERRα, mevalonate pathway, cholesterol, mitochondria

## Abstract

Cellular senescence is a cell program induced by various stresses that leads to a stable proliferation arrest and to a senescence-associated secretory phenotype. Accumulation of senescent cells during age-related diseases participates in these pathologies and regulates healthy lifespan. Recent evidences point out a global dysregulated intracellular metabolism associated to senescence phenotype. Nonetheless, the functional contribution of metabolic homeostasis in regulating senescence is barely understood. In this work, we describe how the mevalonate pathway, an anabolic pathway leading to the endogenous biosynthesis of poly-isoprenoids, such as cholesterol, acts as a positive regulator of cellular senescence in normal human cells. Mechanistically, this mevalonate-induced senescence is partly mediated by the downstream cholesterol biosynthetic pathway. This pathway promotes transcriptional activity of ERRα leading to dysfunctional mitochondria, ROS production, DNA damage and a p53-dependent senescence. Supporting the relevance of these observations, increase of senescence in liver due to a high-fat diet regimen is abrogated in ERRα knockout mouse. Overall, this work unravels the role of cholesterol biosynthesis in the induction of an ERRα-dependent mitochondrial program leading to cellular senescence and related pathological alterations.

## Introduction

Cellular senescence is a program promoted by a myriad of stresses such as telomeres shortening, oncogenic stress-induced hyper-replication and oxidative stress-mediated damages^1^. This program is characterized by a permanent cell cycle arrest and a senescence-associated secretory phenotype (SASP)^1^, both implicated in the pathophysiological effects of senescent cells^2,3^. Senescence-associated pathophysiological contexts include development, tissue regeneration, cancer and aging^2,3^. Although timely regulated senescence exerts beneficial effects, the accumulation of senescent cells throughout life and upon exposure to chronic stresses exerts detrimental effects by promoting aging and its associated diseases^4–9^. Mechanistically, while downstream factors and effectors, such as p53, p21^CIP1^ and p16^INK4A^ or NF-κB and C/EBPβ, respectively blocking cell cycle progression or promoting SASP, were extensively studied^1–3^, upstream molecular and subcellular mechanisms controlling these factors are less understood.

Senescent cells harbour metabolic changes related to both catabolism and anabolism^1,10–15^, to the point that some metabolic specificities of senescent cells are used to target or detect them^11,16 17^. Indeed, senescent cells display metabolic rearrangements as evidenced for instance by an altered glycolytic state and glucose utilization^11,12,18^, a deregulated mitochondrial metabolism, an AMPK activation and an altered NAD^+^ metabolism^10,19–22^. Lipid metabolism is also modified in senescent cells^15,23–30^. For instance, senescent hepatocytes^26,27^, fibroblasts^31^, and T-cells^32^ display an accumulation of lipid droplets (LD), this later accounting mostly for an increase of free fatty acids^23^ and free cholesterol^25,28^, subsequently esterified and incorporated in LD via triglycerides (TG) or cholesteryl esters. Mevalonate (MVA) pathway is part of lipid anabolism and involved in the endogenous biosynthesis of poly-isoprenoids, such as prenyl groups, ubiquinone, cholesterol or dolichol^33^. MVA pathway is thus crucial for many cellular processes including protein-protein interactions, mitochondrial respiration, membranes fluidity or glycosylation^33^. Some studies using pharmacological tools, such as statins or bisphosphonates, have previously suggested an involvement of this pathway in regulating senescence, still with some contradictory effects reported^34,35^. Furthermore, how endogenous cholesterol biosynthesis mechanistically regulate cellular senescence remains so far elusive. Interestingly, in a functional genetic screen of a constitutively active kinase library that we previously reported^36^, two kinases of the MVA pathway, mevalonate kinase (MVK) and phosphomevalonate kinase (PMVK) were identified as able to induce premature senescence when ectopically expressed. Still the mechanisms of action of these enzymes in this context and the relevance of these observations during senescence *in vivo* remained unknown.

In this study, we assessed through the use of genetic tools whether and how the MVA pathway and the downstream biomolecules regulate cellular senescence. Unexpectedly, this work led to the identification of a new mechanism regulating senescence: a cholesterol biosynthetic pathway-dependent ERRα transcriptional program leading to mitochondrial dysregulation and senescence-associated liver alterations.

## Results

### Mevalonate pathway promotes cellular senescence

In order to investigate whether the MVA pathway can regulate senescence, we expressed in MRC5 normal human fibroblasts (i) a constitutively active PMVK^37^ and its kinase dead mutant (PMVKmut)^38^ to examine whether the kinase activity of PMVK is required to induce premature senescence, and (ii) an shRNA directed against PMVK (shPMVK) to examine the role of the MVA pathway during replicative senescence (Fig. 1A-B and Sup. Fig. 1A). Constitutive overexpression of PMVK, but not the kinase dead mutant PMVK, led to decreased cell proliferation (Fig. 1C-D). On the opposite, knocking down PMVK extended the replicative potential of the normal human fibroblasts (Fig. 1C-D). The overexpression of PMVK, but not its kinase dead version, was also associated with the induction of key markers of cellular senescence: senescence-associated β-galactosidase (SA-β-gal) activity (Fig. 1E) and increased mRNA levels of *p21^CIP1^* and SASP marker *IL-8* (Fig. 1F). On the contrary, decreasing PMVK reduced SA-β-gal activity and decreased mRNA levels of *p21^CIP1^* and *IL-8* at late passage (Fig. 1E-F). Together these results show that PMVK contributes to replicative senescence and its kinase activity participates in senescence regulation.

**Figure 1.**
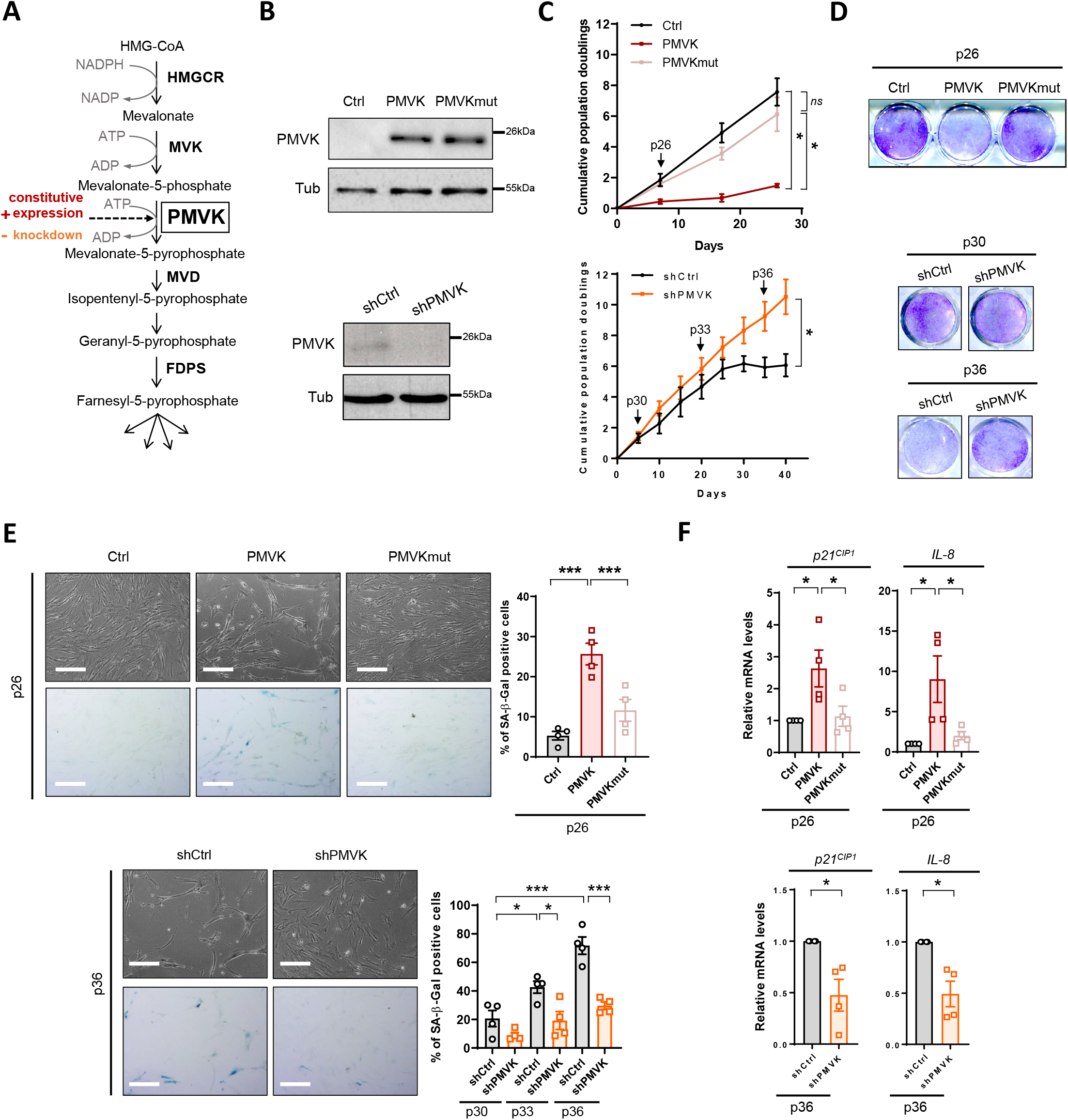
The MVA pathway regulates cellular senescence in normal cells. **A.** Schematic representation of the MVA pathway in eukaryotic cells. HMGCR, HMG-CoA reductase; MVK, mevalonate kinase; PMVK, phosphomevalonate kinase; MVD, mevalonate-diphosphate decarboxylase; FDPS, farnesyl diphosphate synthase. Constitutive expression (in red) or knockdown by shRNA (in orange) was performed in MRC5 fibroblasts. **B.** Relative PMVK and Tubulin protein levels in empty vector-(Ctrl), PMVK- or Kinase Dead (PMVKmut) PMVK-expressing cells and in shCtrl- or shPMVK-expressing cells. **C.** Growth curves of Ctrl, PMVK- or PMVKmut-expressing cells, and shCtrl and shPMVK-expressing cells. Mean +/− SEM of n=4 independent biological replicates. Paired Student’s T-test on last time point. **D.** Crystal violet staining of Ctrl-, PMVK- or PMVKmut-expressing cells at early passage (p26), and of shCtrl and shPMVK-expressing cells at early (p30) and late (p36) passage. **E.** Micrographs and quantification of SA-β-gal positive cells of Ctrl-, PMVK- or PMVKmut-expressing cells at early passage (p26), and of shCtrl and shPMVK-expressing cells during passages. Mean +/− SEM of n=4 independent biological replicates. Scale bar: 10μm. One-way paired ANOVA test. **F.** RT-qPCR of *p21^CIP1^* and *IL-8* genes in Ctrl-, PMVK- or PMVKmut-expressing cells at early passage (p26), and of shCtrl- and shPMVK-expressing cells at late passage (p36). Mean +/− SEM of n=4 independent biological replicates. One-way paired ANOVA test and paired Student’s T-test.

To further prove that this observed senescence is mediated by the MVA pathway and not the sole PMVK enzyme, we stably expressed a shRNA against PMVK and subsequently expressed the upstream enzyme MVK (Fig. 1A). As expected, constitutive overexpression of MVK (Sup. Fig. 1B) induced premature cellular senescence, as shown by decreased cell density (Sup. Fig. 1C-D), elevated SA-β-gal activity (Sup. Fig. 1E) and induction of both *p21^CIP1^* and *IL-8* mRNA levels (Sup. Fig. 1F). Interestingly the knockdown of PMVK abolished MVK-induced premature senescence, rescuing decreased cell density (Sup. Fig. 1C-D), and increased SA-β-gal activity and *p21^CIP1^* and *IL-8* mRNA levels (Sup. Fig. 1E-F).

Taken together, these data support that the MVA pathway plays a pivotal role in promoting cellular senescence.

### Dysfunctional mitochondria, ROS production, DNA damage and p53 mediate mevalonate pathway-induced senescence

P53 is a well-known critical effector of cellular senescence^39^. Its transcriptional targets such as *p21^CIP1^*, *GADD45A* and *GDF15* were upregulated at mRNA levels by the constitutive expression of PMVK (Fig. 2A) suggesting that p53 could mediate PMVK-induced senescence. Indeed, p53 knockdown by siRNA in PMVK overexpressing cells (Sup. Fig. 2A) reverted PMVK-induced p53 targets and *IL-8* mRNA expression (Fig. 2A), decrease in cell number (Fig. 2B) and increase in SA-β-Gal activity (Fig. 2C).

**Figure 2.**
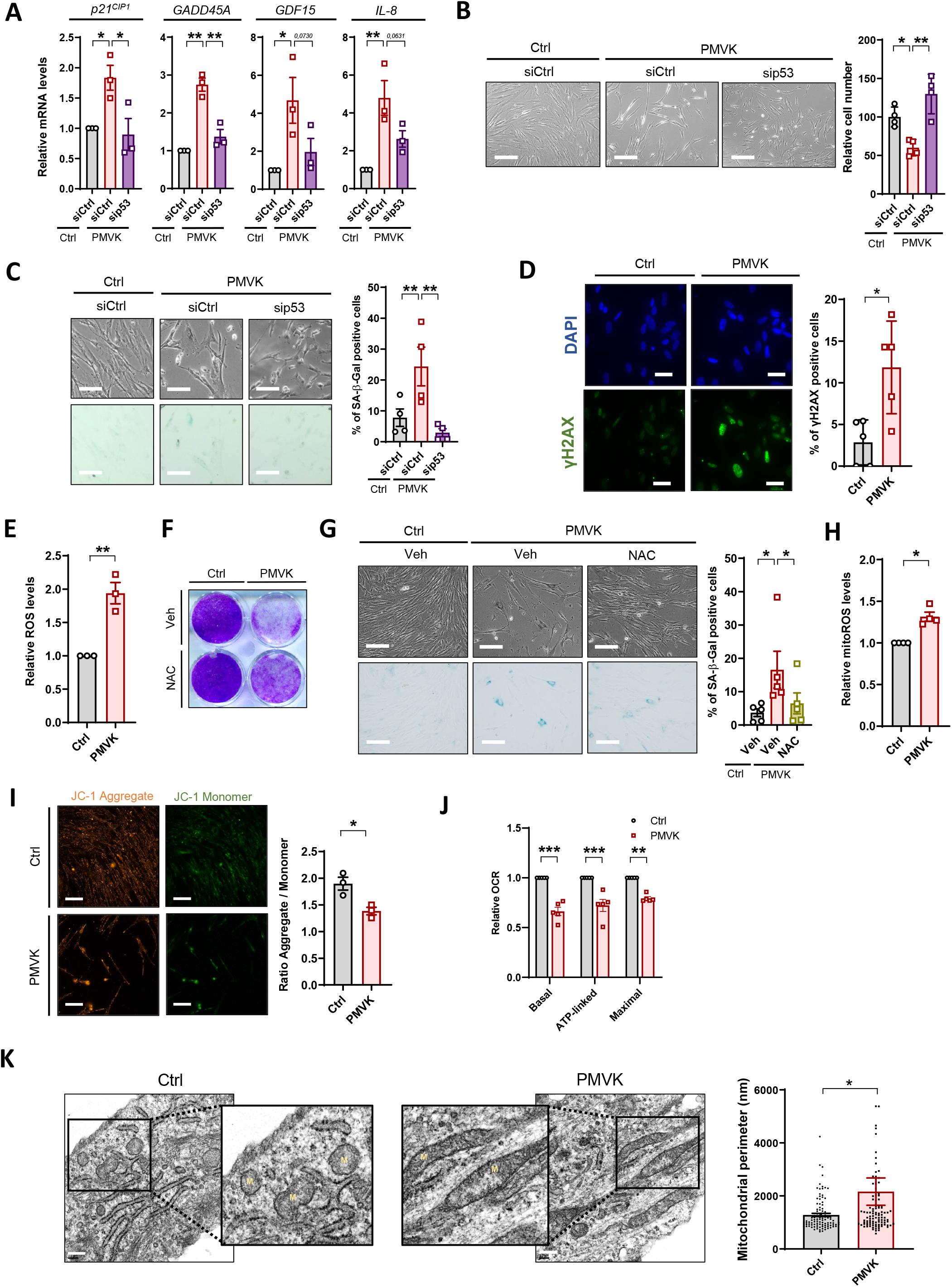
MVA pathway induction promotes mitochondrial dysfunction, ROS accumulation, DNA damage and p53-dependent senescence. **A.** RT-qPCR of p53 target-genes, including *p21^CIP1^, GADD45A, GDF15* and *IL-8* in Ctrl or PMVK-expressing cells previously transfected with control non-targeting siRNA (siCtrl) or p53-targeted siRNA (sip53). Mean +/− SEM of n=3 independent biological replicates. One-way paired ANOVA test. **B.** Micrographs of Ctrl and PMVK-expressing cells upon sip53. Scale bar: 10μm. Quantification of relative cell number in Ctrl and PMVK-expressing cells. Mean +/− SEM of n=4 independent biological replicates. Scale bar: 10μm. One-way paired ANOVA test. **C.** Representative micrographs and quantification of SA-β-gal positive cells in Ctrl and PMVK-expressing cells upon siCtrl or sip53. Mean +/− SEM of n=4 independent biological replicates. Scale bar: 5μm. One-way paired ANOVA test. **D.** Micrographs of Ctrl or PMVK-expressing cells immunostained with γH2AX antibody and DAPI stained. Scale bar: 5μm. Quantification of γH2AX positive cells in Ctrl and PMVK-expressing cells. Mean +/− SD of one representative experiment (n=2). Unpaired Student’s T-test. **E.** Quantification of total ROS in Ctrl and PMVK-expressing cells. Mean +/− SEM of n=3 independent biological replicates. Paired Student’s T-test. **F.** Crystal violet staining after 8 days in Ctrl and PMVK-expressing cells upon vehicle (Veh) or N-acetyl-cysteine (NAC) treatment. **G**. Representative micrographs and quantification of SA-β-gal positive cells in Ctrl and PMVK-expressing cells upon vehicle or NAC treatment. Mean +/− SEM of n=5 biological replicates. Scale bar: 10μm. One-way paired ANOVA test. **H.** Quantification of mitochondrial ROS in Ctrl and PMVK-expressing cells. Mean +/− SEM of n=4 independent biological replicates. Paired Student’s T-test. **I.** Mitochondrial membrane polarisation measurement assessed by JC1-probe. Representative micrographs of JC1 in both monomer (green) or aggregate (orange) forms within Ctrl and PMVK-expressing cells. Scale bar: 10μm. Quantification of ratio fluorescence intensity (Monomer) / fluorescence intensity (Aggregate). Mean +/− SEM of n=3 independent biological replicates. Paired Student’s T-test. **J.** Quantification of relative oxygen consumption rate (OCR) assessed thanks to Seahorse analysis. Mean +/− SEM of n=5 independent biological replicates. Multiple paired Student’s T-tests. **K.** Representative electron micrographs of mitochondria in Ctrl and PMVK-expressing cells and quantification of mitochondrial perimeters. M: Mitochondria. Mean +/− SD of n=103 (Ctrl) and n=106 (PMVK) mitochondria. Representative of n=3 independent experiments. Scale bar: 300nm.

During cellular senescence, p53 activation can occur through oxidative stress-induced DNA damage ^1^ and its subsequent DNA damage response^2,3^. Accordingly, p53 pathway activation upon PMVK overexpression resulted from increased oxidative stress - DNA damage pathway as PMVK overexpressing cells displayed increased DNA damage as shown by increased γH2AX positive cells (Fig. 2D), concomitantly with increased total ROS levels (Fig. 2E). Importantly, NAC anti-oxidant treatment largely overcame PMVK-induced senescence (Fig. 2F-G) showing a critical role of ROS in mediating MVA-induced senescence.

Mitochondria is one of the major sites of ROS production in the cell^40^ and its dysregulation participates in cellular senescence^13,19,41^. Strikingly, constitutive overexpression of PMVK led to various mitochondrial functional and morphological alterations: mitochondrial ROS generation (Fig. 2H), mitochondrial depolarization (Fig. 2I), decreased respiration (Fig. 2J) and increased mitochondrial size (Fig. 2K).

These results support that PMVK promotes senescence by dysregulating the mitochondria, inducing ROS production, DNA damage and p53 activation.

### The cholesterol biosynthetic branch participates in mevalonate pathway-induced senescence

Farnesyl-5-pyrophosphate is the end-product of the MVA pathway and presents three isoprene units, elementary units further used either to be transferred as prenyl groups to proteins or to be condensated into more complex poly-isoprenoids, that include among others dolichol, ubiquinone or cholesterol^33^ (Fig. 3A). In order to dissect the role of downstream branches in the MVA-induced senescence, we knocked down the first enzyme of each branch, subsequently overexpressed PMVK and automatically counted the number of cells 4 days later. Only the knockdown of FDFT1, the first enzyme of the cholesterol biosynthesis branch, partially reverted the decreased cell number induced by PMVK overexpression, an effect comparable to the one observed with knockdown of FDPS, the last enzyme of the MVA pathway (Fig. 3A-B). SiRNA-mediated FDFT1 knockdown (Sup. Fig. 2B) also rescued PMVK-induced decrease in cell density (Fig. 3C), increase in SA-β-gal activity (Fig. 3D), and increase in *p21^CIP1^* and *IL-8* mRNA levels (Fig. 3E), without impacting *PMVK* mRNA level (Sup. Fig. 2C). Confirming these results, stable knockdown of FDFT1 by two independent shRNA largely reverted premature senescence induced by the MVA pathway (Sup. Fig. 2D-F).

**Figure 3.**
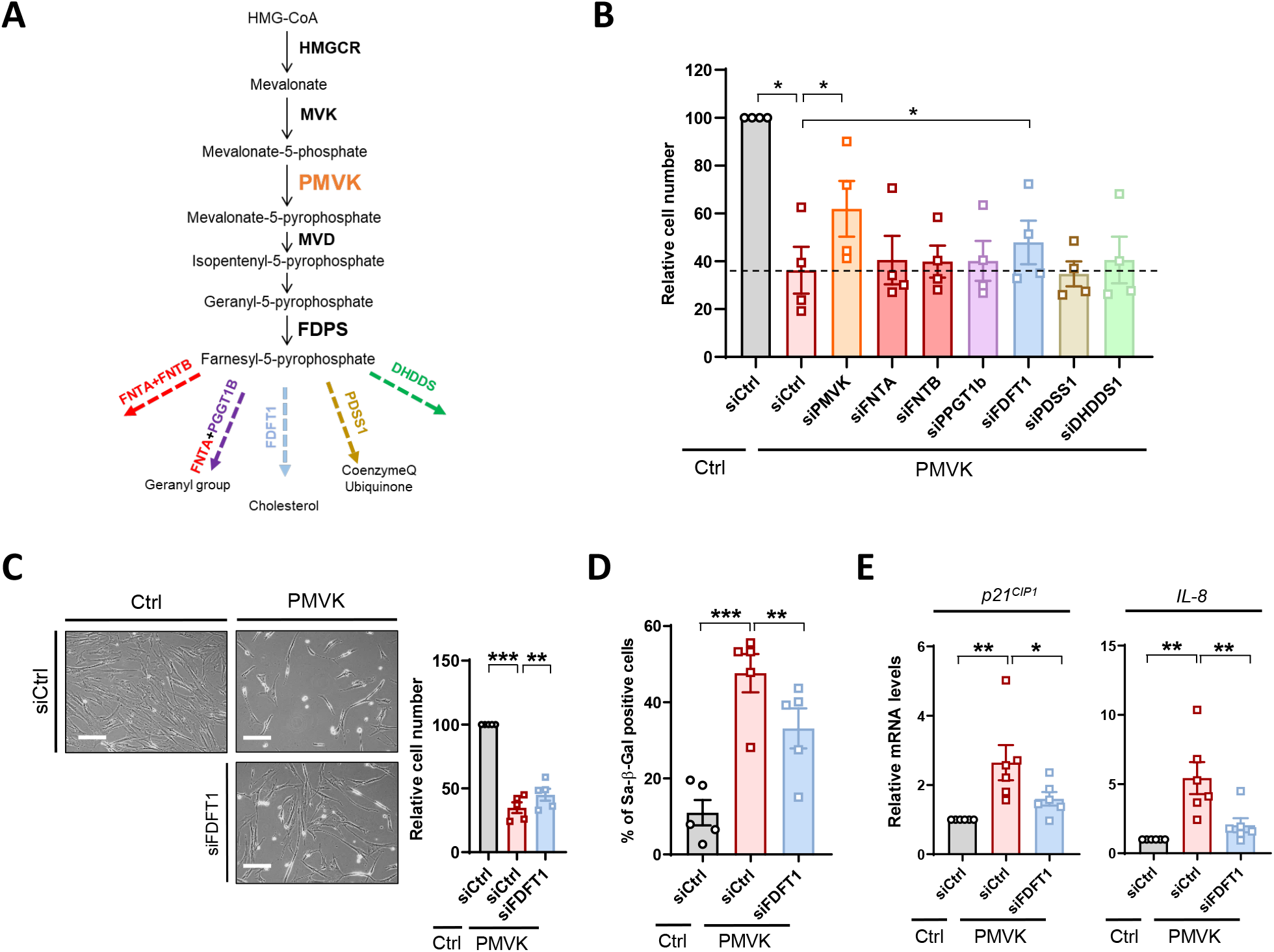
Cholesterol biosynthetic branch participates in MVA-induced senescence. **A.** Schematic representation of the MVA pathway and subbranches stemming from Farnesyl-5-Pyrophosphate, including farnesylation (red), geranylation (purple), cholesterol synthesis (blue), ubiquinone synthesis (gold) and dolichol biosynthesis (green). First specific enzymes of each subbranch are indicated. **B.** Quantification of number of cells after transfection with siRNA against PMVK and enzymes of downstream subbranches of the MVA pathway in PMVK-expressing cells. Mean +/− SEM representative of n=4 independent biological replicates. One-way paired ANOVA test. **C.** Representative micrographs of Ctrl and PMVK-expressing cells upon siCtrl or siFDFT1 transfection (scale bar: 20μm) and cell number quantification (mean +/− SEM of n=5 independent biological replicates, One-way paired ANOVA test). **D.** Quantification of SA-β-gal positive cells in Ctrl and PMVK-expressing cells upon siCtrl or siFDFT1 transfection. Mean +/− SEM of n=5 independent biological replicates. One-way paired ANOVA test**. E.** RT-qPCR of *p21^CIP1^* and *IL-8* genes in Ctrl and PMVK-expressing cells upon siCtrl or siFDFT1 transfection. Mean +/− SEM of n=6 independent biological replicates. One-way paired ANOVA test.

Altogether these results indicate a role of the cholesterol biosynthesis pathway in the regulation of mevalonate-induced senescence.

### Mitochondrial master regulator Estrogen-Related Receptor alpha mediates mevalonate-induced senescence

Estrogen-Related Receptor alpha (ERRα), which has been proposed to be activated by cholesterol, is a key regulator of mitochondrial functions and of metabolism^42,43^. We then assessed during PMVK-induced senescence the expression of ERRα targets: ERRα itself^44^ encoded by *ESRRA* gene and *UQCRFS1*, *NDUF5A*, *SDHA* and *SDHB* nuclear-encoded mitochondrial targets^42^. The constitutive overexpression of PMVK increased the expression of *ESRRA* (Fig. 4A and Sup. Fig. 3A), *UQCRFS1*, *NDUF5A*, *SDHA* and *SDHB* (Fig. 4B and Sup. Fig. 3A) and this upregulation was largely reverted by the knockdown of FDFT1 (Fig. 4A-B and Sup. Fig. 3A) supporting the hypothesis of ERRα activation by the cholesterol biosynthetic pathway in this context. Further confirming an activation of ERRα during MVA-induced senescence, its knockdown by siRNA strategy or its activity inhibition using XCT-790 pharmacological inhibitor^45^ blocked MVA-induced upregulation of various ERRα targets (Fig. 4C-D and Sup. Fig. 3B), without impacting PMVK levels (Sup. Fig. 3C).

**Figure 4.**
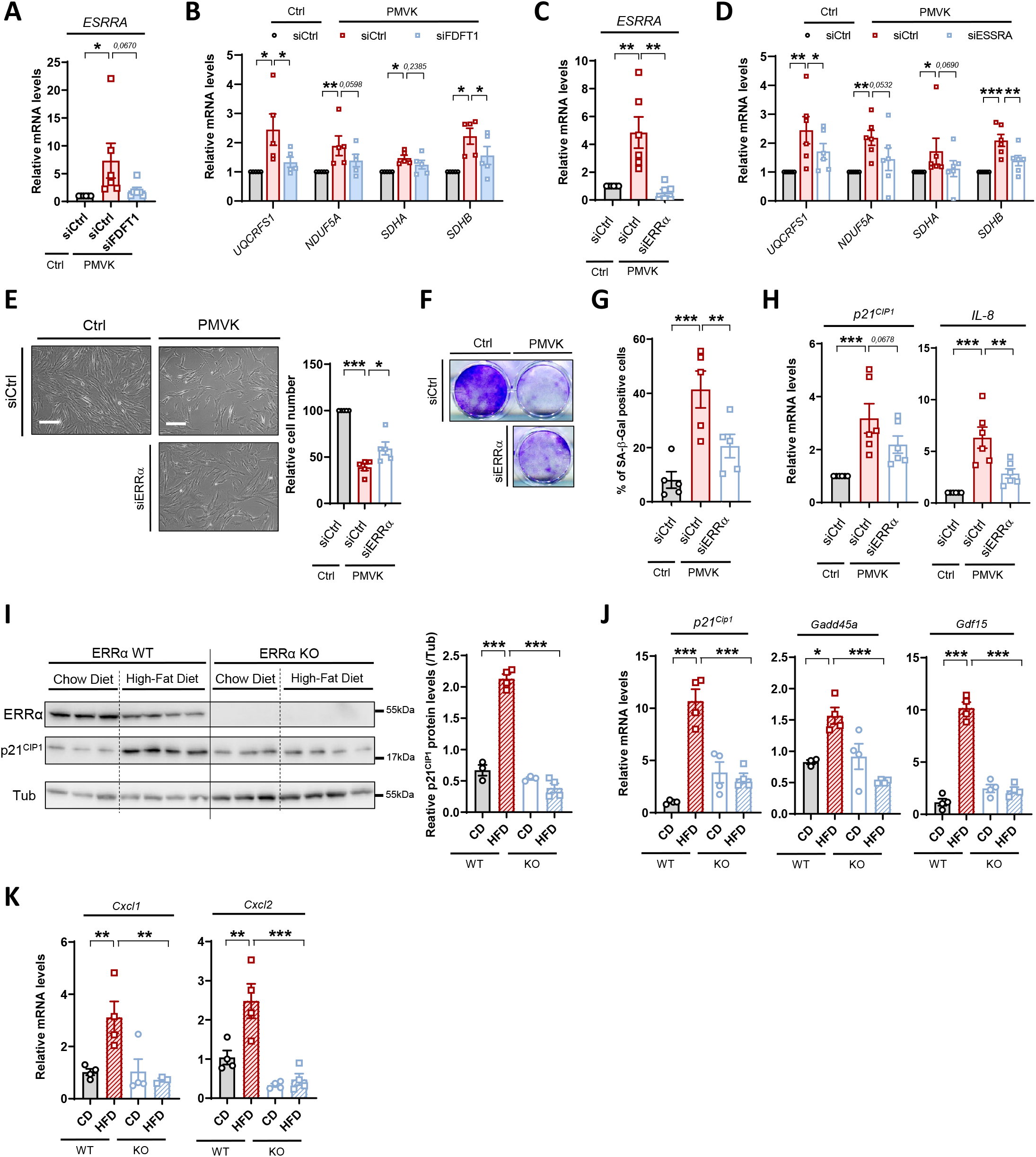
Mitochondrial master regulator Estrogen-Related Receptor alpha mediates mevalonate-induced senescence. **A-B.** RT-qPCR of *ESSRA* and ERRα target genes (including *UQCRSF1, NDUF5A, SDHA, SDHB*) in Ctrl and PMVK-expressing cells, upon siCtrl or siFDFT1 transfection. Mean +/− SEM of n=5-6 independent biological replicates. One-way paired ANOVA test. **C-D.** RT-qPCR of *ESRRA* and ERRα target genes in Ctrl and PMVK-expressing cells, upon siCtrl or siERRα transfection. Mean +/− SEM of n=6 independent biological replicates. One-way paired ANOVA test. **E.** Representative micrographs and cell number quantification of Ctrl and PMVK-expressing cells previously transfected with siERRα. Scale bar: 20μm. Mean +/− SEM of n=5 independent biological replicates. One-way paired ANOVA test. **F.** Crystal violet staining of Ctrl and PMVK-expressing cells previously transfected with siERRα. **G.** Quantification of SA-β-gal positive cells in Ctrl and PMVK-expressing cells upon siERRα transfection. Mean +/− SEM of n=5 independent biological replicates. One-way paired ANOVA test. **H.** RT-qPCR of *p2i^CIP1^* and *IL-8* genes in Ctrl and PMVK-expressing cells previously transfected with siCtrl or siERRα. Mean +/− SEM of n=6 independent biological replicates. One-way paired ANOVA test. **I.** Western blot on ERRα, p21^CIP1^ and Tubulin in liver of ERRα WT and ERRα KO mice fed either by chow diet (CD) or high-fat diet (HFD). Quantification of p21^CIP1^ levels normalized to Tubulin levels. Mean +/− SEM of n=3-4 mice. One-way paired ANOVA test. **J-K.** RT-qPCR of p53 target-genes (namely *p21^Cipl^*, *Gadd45a*, *Gdfl5*) and pro-inflammatory *Cxcl1* and *Cxcl2* genes in liver of ERRα WT and ERRα KO mice fed either by chow diet (CD) or high-fat diet (HFD). Mean +/− SEM of n=3-4 mice. One-way paired ANOVA test.

We next sought to determine whether cholesterol-dependent ERRα program functionally mediates the MVA-induced senescence. Knockdown of ERRα in PMVK-overexpressing cells partially rescued the decreased cell number (Fig. 4E-F), the increased SA-β-gal activity (Fig. 4G), and the elevated *p21^CIP1^* and *IL-8* mRNA levels (Fig. 4H). Further supporting that ERRα mediates MVA/cholesterol biosynthetic pathway-induced senescence, its chemical inhibition using XCT-790 compound, similarly to ERRα knockdown, largely bypassed senescence induced by PMVK expression (Sup. Fig. 3D-G).

To next investigate the relevance of these discoveries in a relevant pathophysiological system, we investigated the level of cellular senescence in the liver of ERRα knockout mice fed by high fat diet (HFD), known to increase cholesterol levels and drive cellular senescence and senescence-dependent steatosis^27,46^. ERRα target genes *Uqcrfs1*, *Nduf5a*, *Sdha* and *Sdhb* were accordingly reduced at mRNA levels in liver of ERRα knockout mice (Sup. Fig. 4H). As expected, HFD resulted in accumulation of senescence markers such as *p21^Cip1^* (Fig. 4I-J), *Gadd45a* and *Gdf15* p53 targets (Fig. 4J), and pro-inflammatory SASP factors *Cxcl1* and *Cxcl2* (Fig. 4K), which are murine functional orthologs of human *IL8*^47^. Strikingly, increase in the expression of these senescence markers was abrogated in liver of ERRα knockout mice upon HFD (Fig. 4I-J-K). In addition and as previously reported^48^, HFD-fed ERRα knockout mice did not display hepatic steatosis (Sup. Fig. 4I), a process tightly linked to senescent cell accumulation^27,46^.

Overall, these results highlight the importance of ERRα in mediating MVA/cholesterol biosynthetic pathway-induced cellular senescence *in vitro* in human cells and *in vivo* in mouse.

## Discussion

In this study, we deciphered the role of the MVA pathway, the cholesterol biosynthetic pathway and of ERRα transcription factor in mediating cellular senescence. MVA pathway activation triggers premature senescence whereas its inhibition delays replicative senescence in normal human cells. MVA-induced senescence is mediated, at least partly, by the biosynthetic cholesterol pathway and the activation of an ERRα transcriptional program, mitochondrial ROS accumulation, DNA damage and p53 activation.

Several studies depicted a role of MVA pathway in cellular senescence through the use of statins and aminobisphosphonates inhibitors but with conflicting conclusions. For instance blocking the MVA pathway with these inhibitors has been shown to delay senescence in HUVEC model^34^, whereas it promotes senescence in oral keratinocytes and has no obvious effect in oral fibroblasts^35^. Besides, statins and aminobiphosphonates have been shown both to blunt SASP, notably pro-inflammatory cytokines including IL-6, IL-8 and monocyte chemoattractant protein (MCP)-1^49–52^, and some of its effects, including pro-tumorigenic activity^53^. Dampening SASP through statins could explain their numerous anti-inflammatory outcomes *in vivo*^54,55^. Our results using genetic tools, knockdown and/or overexpression of PMVK or MVK, clearly demonstrate that the MVA pathway promotes cellular senescence and associated SASP in normal human fibroblasts. Discrepancies with pharmacological studies might thus result from the known pleiotropic effects of the MVA inhibitors, as these inhibitors were found to exert multiple MVA-independent actions^56,57^.

Our results reveal an unexpected link between the MVA pathway and the DNA damage/p53 pathway, a well-known effector pathway promoting cellular senescence^58^. This DNA damage/p53 pathway was found to be activated by an oxidative stress upon MVA-induced senescence, probably mainly provoked by increased mitochondrial ROS production.

Mitochondrial ROS are well known mediators of cellular senescence and can result from multiple mitochondrial alterations, from increased respiration and thus increased basal electron leaks during the electron transport chain activity, to decreased respiration but increased levels of electron leaks because of altered electron transport chain complex associations^40^. According to our findings, MVA-induced senescence results in decreased respiration with increased mitochondrial ROS production. This is in line with previous results showing that maladaptive complex 1 assembly through increased transcriptional program of ETC components results in decreased respiration with increased mitochondrial ROS and promotes cellular senescence and aging^59^.

Our siRNA approach to reveal which sub-branch(es) of the MVA pathway participate in MVA-induced senescence identified an involvement of the cholesterol biosynthetic branch. Of note, this finding does not exclude that some other sub-branches could collaborate with the cholesterol biosynthetic pathway to regulate cellular senescence. Beyond the specific role of ceramide^24^, little is known about lipids and senescence except that the lipid metabolism is largely modified in senescent cells. In particular it has been reported that free fatty acids and cholesterol levels are increased in senescent cells^25,26,28,29^ and that ceramides or prostaglandins regulate also the onset of senescence^24,60^. In line with our findings, a recent article reported an upregulation of the expression of cholesterol synthesis genes in a model of oncogene-induced senescence^61^. Noteworthy, knockdown of some cholesterol synthesis genes partially bypassed cell proliferation arrest in this system. However, the mechanisms behind these observations were not deciphered^61^. Our results underline the importance of cholesterol biosynthetic branch downstream of the MVA pathway in the regulation of cellular senescence and allow us to propose a mechanistic model through ERRα activation.

Our results support that ERRα mediates the pro-senescence function of the cholesterol biosynthetic pathway as its knockdown or its chemical inhibition overcomes MVA-induced premature senescence. A previous study proposed that cholesterol can be an ERRα ligand^43^, as this is still largely debated further investigation will be needed to confirm or not this observation. As far as we know ERRα has not been associated with cellular senescence. Nevertheless, it has been shown to be activated during cancer cell death in response to PLA2R1/JAK/STAT signalling, this signalling being known to promote cellular senescence in normal human cells and in mice^51,62–64^. In cancer cells, PLA2R1/JAK/STAT signalling in an ERRα-dependent manner blocks mitochondrial respiration, alters mitochondria and mediates ROS generation^65^, reminiscent to the observed alterations during MVA-induced senescence in normal cell.

We have previously reported that ERRα-null mice are protected from HFD-induced non-alcoholic fatty liver disease (NAFLD)^48^, which is characterized by liver steatosis. Senescent cells accumulate in the liver during HFD and/or aging and contribute to liver steatosis^27,46^. We observed that ERRα-depleted livers are protected from increased senescence and steatosis, suggesting that reduced ERRα-induced senescence could contribute to this decreased steatosis in the liver of these mice. Mechanistically, it has been previously shown that ERRα loss^48^ or elimination of senescent cells^27^ results in change in free fatty acid metabolism decreasing liver steatosis. Beyond these observations highlighting the importance of ERRα in regulating senescence and senescence-dependent liver steatosis, we can speculate that other known pathophysiological processes regulated by ERRα could rely on its effect on cellular senescence. For instance, ERRα-null mice display reduced osteoporosis in female mice^66,67^ and cellular senescence in the bone is known to promote osteoporosis^68^, suggesting that ERRα-promoted senescence could participate in ERRα-induced osteoporosis. In the same line, anti-osteoporosis effects of MVA pathway inhibitors, aminobisphosphonates, are well known^69^ and may partly rely on reduced ERRα activity, though further studies need to critically test this hypothesis.

Overall, our results emphasize a MVA/cholesterol biosynthetic pathway/ERRα program as a new pathway regulating cellular senescence. As impacting cellular senescence has attracted many interests to improve various age-related diseases and health span^7-9^, our work paves the way to a new potential senotherapeutic strategy using drugs targeting ERRα.

## Methods

### Cell culture and reagents

MRC5 normal human fibroblasts (ATCC, Manassas, VA, USA), and kidney 293 GP cells (Clontech, Mountain View, CA, USA) were cultured in Dulbecco’s modified Eagle’s medium (DMEM, Life Technologies, Carlsbad, USA) with GlutaMax and supplemented with 10% FBS (Sigma-Aldrich, Saint-Louis, USA) and 1% penicillin/streptomycin (ThermoFisher Scientific). XCT-790 (Medchem, HY-10426/CS-2413) was used at 150 nM. N-Acetyl-Cysteine (NAC) (A9165, Sigma-Aldrich) was used directly after infection at 1 mM. These compounds were renewed every two days.

### Vectors, transfection and infection

Retroviral vectors were used to constitutively overexpress MVK or PMVK. Vector plasmids were provided by Addgene in the Myristoylated Kinase Library (Kit #1000000012) described in^37^. K22M mutation of the kinase dead PMVK mutant (PMVKmut) was generated using QuickChange Site-Directed Mutagenesis Kit (Agilent, Catalog # 200518). Lentiviral particles were used to constitutively express shPMVK (pLV[shRNA]-Hygro-U6> hPMVK[shRNA#4], Target sequence: GAGAACCTGATAGAATTTATC) and shFDFT1 (pLV[shRNA]-Hygro-U6> hFDFT1[shRNA#2], Target sequence CAACGATCTC CCTTGAGTTTA and pLV[shRNA]-Hygro-U6> hFDFT1[shRNA#3], Target sequence: ACC ATTTGAATGTTCGTAATA) and were provided by VectorBuilder. 293T or 293GP virus producing cells were transfected using the GeneJuice reagent according to the manufacturer’s recommendations (Merck Millipore). Two days after transfection, viral supernatant was collected, diluted with fresh medium (1/2) and hexadimethrine bromide was added (final concentration 8 μg/ml; Sigma-Aldrich). Target cells were then infected, centrifugated with virus particles for 30 minutes at 2000 rpm and subsequently incubated during 6 hours at 37°C 5% CO2. Fresh medium was added after 6 hours incubation. One day later, infected cells were selected with Geneticin (ThermoFisher Scientific) at 75 μg/mL.

### siRNA

MRC5 fibroblasts were plated and reverse transfected with ON-TARGET plus SMART pool of small interference (si) RNAs: siCtrl (Catalog: D-001810-10-20 / Lot#2693147), siPMVK (Catalog: L-006782-00), siFDPS (Catalog: L-008632-00), siFNTA (Catalog: L-008807-00), siFNTB (Catalog: L-010093-00), siPGGT1B (Catalog: L-008703-00), siFDFT1 (Catalog: #L-009442-00-0005 / Lot#180518), siPDSS1 (Catalog: L-008464-01), siDHDDS (L-010399-01), siESRRA (Catalog: #L-003403-00-0005 / Lot#191211), sip53 (Catalog: #L-003329-00-0001) / Lot#180912) (Horizon Discovery) previously incubated for 20 min with Dharmafect 1 Transfection Reagent (Horizon Discovery) 0,6% in antibiotics- and serum-free medium DMEM with Glutamax. Final siRNA concentration in the well was 15nM. The day after, cells were infected (with retroviral particles containing PMVK) as referenced previously.

### Animals

Wild-type (WT) and ERRα^-/-^ mice in a C57BL/6J genetic background were housed and fed ad libitum with free access to water in an animal facility at McGill University. All animal experiments were conducted in accord with accepted standards of humane animal care and all protocols were approved by the McGill Facility Animal Care Committee and the Canadian Council on Animal Care. For high-fat diet (HFD) experiments, mice were separated randomly into groups of two to three mice per cage and fed either a control chow diet consisting of 10 kcal percent fat (catalog no. TD.08806; Harlan, Indianapolis, IN) or an HFD consisting of 60 kcal percent fat (catalog no. TD.06414; Harlan) during 15 weeks, initiated at 6 weeks of age. For all mouse experiments, littermates were used and mice were euthanized by cervical dislocation at Zeitgeber time (ZT) 4 for tissue isolations.

### RNA extraction, reverse transcription, and real-time quantitative PCR

Total RNAs were extracted with phenol-chloroform using Upzol (Dutscher, Brumath, France). Synthesis of cDNA was performed using Maxima First cDNA Synthesis Kit (ThermoFisher Scientific) from 1 μg of total RNA. cDNA (50 ng/μL) was used as a template for quantitative PCR (qPCR), and mixed with primers (200 nM), SYBR^™^ Green PCR Master Mix (ThermoFisher Scientific) or TaqMan mix (Roche) and Universal Probe Library probes (100 μM) (ThermoFisher Scientific) for the gene of interest. Reactions were performed in triplicate. qPCR analyses were carried out with the FX96 Thermocycler (Biorad, Hercules, USA). Relative mRNA levels were calculated using the Comparative Ct (ΔΔCT) method. Gene expression was normalized with *hACTB* or *mRplp*. Primer sequences used are listed in Supplementary Table S1.

### Senescence-associated β-Galactosidase analysis and Crystal violet

For SA-β-Galactosidase assay, cells were washed with PBS 1X, fixed for 5 min in 2% formaldehyde / 0.2% glutaraldehyde, rinsed twice in PBS 1X, and incubated at 37°C overnight in SA-β-Galactosidase staining solution as previously described^70^. For crystal violet staining, cells were washed with PBS 1X, fixed for 15 min in 3.7% formaldehyde and stained with crystal violet.

### ROS and JC1 quantification

Total cellular and specific mitochondrial ROS were measured respectively with CellROX™ Green Reagent (ThermoFisher Scientific) and Cell Meter™ Mitochondrial Hydroxyl Radical Detection Kit (ATT Bioquest) according to manufacturer’s recommandations. For JC1, JC1-Mitochondrial Membrane Potential Assay Kit (ab113850, Abcam) was used. JC1 monomers and aggregates were both excited at 488 nm. Detection of fluorescence for JC1 monomers and aggregates were performed respectively at 530nm and 590nm. Ratio F(aggregate)/F(monomer) was subsequently evaluated. Pictures acquisition was performed using Operetta CLS High-Content Analysis System (PerkinElmer). All the quantifications of ROS and JC1 were performed using Columbus Software.

### Lipid droplets measurements

To assess hepatic steatosis, livers were fixed in 10% buffered formalin and blocked in paraffin. Paraffin-embedded tissues were sectioned (4 mm) and stained with Oil Red O. In brief, sections were washed with distilled water and then stained with 0.5% Oil Red O in propylene glycol for 16 hours. Subsequently, sections were incubated for 1 minute in 85% propylene glycol and washed twice with water. Finally, slides were counterstained with hematoxylin.

### Seahorse

Cells were plated in Seahorse seeding 24-well plates prior infection. Infection was performed in the 24-well plates and cells were subsequently selected for 5 days with Geneticin (ThermoFisher Scientific) at 75 μg/mL. Using Seahorse XF Cell Mito Stress Test (Agilent, Santa Clara), fibroblasts were sequentially treated with 1,5 μM oligomycin, 1,5 μM phenylhydrazone (FCCP), and 0.5 μM Rotenone and Antimycin A. After assay, cells were stained with Hoescht for 10mn before proceeding to cell number counting and subsequent normalization. Data were obtained at SFR Biosciences AniRA-ImmO Platform, ENS de Lyon. Seahorse XFe Wave Software (Agilent) was used to subsequently analyze the data.

### Electron microscopy

1:1 volume of glutaraldehyde 4% was added to the culture medium and cells were incubated 15 min at 4 °C. After discarding medium/glutaraldehyde, 1:1 volume of glutaraldehyde 4 % / cacodylate 0.2 M pH 7.4 was added. After fixation in glutaraldehyde 2%, cells were washed three times for 1 hr at 4°C, post-fixed with 2% OsO4 1 hr at 4°C, and dehydrated with an increasing ethanol gradient. Impregnation was performed with Epon A (50%) plus Epon B (50%) plus DMP30 (1.7%). Inclusion was obtained by polymerisation at 60°C for 72 hr. Ultrathin sections (approximately 70 nm thick) were cut on a UCT (Leica) ultramicrotome, mounted on 200 mesh copper grids and contrasted with uranyl acetate and lead citrate. Acquisition of at least 100 mitochondria of 10-20 independent cells per condition was performed with a Jeol 1400JEM (Tokyo, Japan) transmission electron microscope equipped with an Orius 600 camera and Digital Micrograph at CIQLE platform (UCBL-Lyon).

### Immunoblot and Immunofluorescence

For immunoblot experiments, cells were lysed in RIPA buffer. After protein quantification using Bradford assay, 30 μg of proteins were loaded and resolved by SDS-PAGE electrophoresis and transferred to nitrocellulose membranes (Bio-Rad). Membranes were blocked with TBS Tween / Milk 5% for 1 hour and incubated at 4°C with primary antibodies overnight. Membranes were then incubated with secondary antibody for 1 hour at room temperature. Detection was performed using ECL kit (Amersham). For immunofluorescence experiments, cells were washed with PBS 1X, fixed for 15 minutes in 3.7% formaldehyde and permeabilized with Triton 100X 0,1% for 10 minutes. Blocking was performed using PBS with 20% FBS during 30 minutes and cells were then incubated at 4°C with primary antibodies overnight. Cells were incubated with secondary antibody for 1 hour at room temperature, and washed before proceeding to image acquisition and analyses. Quantification was performed with ImageJ software. All primary antibodies and dilutions used are listed in Supplementary Table S2.

### Statistical analysis

Values represent mean ± SD or SEM as indicated in the figure legend. For *in vivo* experiments, two-sided Grubb’s test was performed to find outliers, which were removed if *p-value* was below 0.05. Statistical analyses for groups were performed as indicated in the figure legend. D’agostini & Pearson normality test was used before proceeding to any analyses. Parametric tests were two tailed, unpaired or paired: Student’s t test (equal variance) or Welch’s t-test (for non-equal variance). Mann-Whitney U Test was performed for non-parametric tests. Spearman test was used for correlations analysis. All the statistical analyses were performed using GraphPad Prism 7 (* P < 0.05; ** P < 0.01; *** P < 0.001).

## Author contributions

D.V.Z., J.C.H. and C.M. performed *in vitro* experiments. M.V., C.S. and J.R. performed *in vivo* experiments. D.V.Z., M.V., V.G., N.M. and D.B. designed the experiments and the results were analyzed by all the co-authors. D.V.Z., N.M. and D.B. designed the overall study. D.B. and N.M. co-supervised the work. D.V.Z., N.M. and D.B. wrote the manuscript with input from all authors.

## Acknowledgements

We thank laboratory members for helpful suggestions and collaborations. This work was supported by the Ligue Régionale contre le Cancer (Comité du Rhône) (NM) and ANR (ANR-14-CE12-0003) (DB). DZ was supported by the French Ministry of Higher Education and Research and by the Fondation pour la Recherche Médicale FRM (FDT201904008259).

