## Supplemental Figures and Tables for "Cholesterol biosynthetic pathway induces cellular senescence through ERRα"

**Figure 1.**

**A.** RT-qPCR of PMVK in Ctrl-, PMVK- or PMVKmut-expressing cells, and shCtrl or shPMVK. Mean +/- SEM of n=4 independent biological replicates. One-way paired ANOVA test and paired Student's T-test. **B.** Relative MVK and Tubulin protein levels in empty vector-(Ctrl), and MVK-expressing cells (MVK). **C.** Representative micrographs (scale bar: 20µm) and cell number quantification (mean +/- SEM of n=4 independent biological replicates. One-way paired ANOVA test) of shCtrl- and shPMVK-expressing cells 5 days after expressing empty vector (Ctrl) or MVK. **D.** Crystal violet staining after 8 days of MVK expression in shCtrl- and shPMVK-expressing cells. **E.** Quantification of SA-β-gal positive cells after 5 days of MVK expression in shCtrl- and shPMVK-expressing cells. Mean +/- SEM of n=4 independent biological replicates. One-way paired ANOVA test **F.** RT-qPCR of *PMVK*, *MVK*, $p21^{CIP1}$ and *IL-8* genes after 5 days of MVK expression in shCtrl- and shPMVK-expressing cells. Mean +/- SEM of n=3 independent biological replicates. One-way paired ANOVA test.

**Figure 2.**

**A.** RT-qPCR of *TP53* gene in Ctrl or PMVK-expressing cells previously transfected with control non-targeting siRNA (siCtrl) or p53-targeted siRNA (sip53). Mean +/- SEM of n=3 independent biological replicates. One-way paired ANOVA test. **B.** RT-qPCR of *FDFT1* gene in Ctrl or PMVK-expressing cells previously transfected with control non-targeting siRNA (siCtrl) or FDFT1-targeted siRNA (siFDFT1). Mean +/- SEM of n=6 independent biological replicates. One-way paired ANOVA test. **C.** RT-qPCR of *PMVK* gene in cells constitutively expressing PMVK, and previously transfected with siCtrl or siFDFT1. Mean +/- SEM of n=3 independent biological replicates. One-way paired ANOVA test. **D.** Crystal violet staining at day 12 after constitutive expression of PMVK in shCtrl, shFDFT1#2 and shFDFT1#3-expressing cells. **E.** Representative micrographs and quantification of SA-β-galactosidase positive cells after constitutive expression of PMVK in shCtrl, shFDFT1#2 and shFDFT1#3-expressing cells. Mean +/- SEM of n=3 independent biological replicates. One-way paired ANOVA test. **F.** RT-qPCR of *FDFT1*, $p21^{CIP1}$ and *IL-8* genes after constitutive expression of PMVK in shCtrl or shFDFT1#2 and shFDFT1#3 cells. Mean +/- SEM of n=3 independent biological replicates. One-way paired ANOVA test.

**Figure 3.**

**A.** RT-qPCR of *ERRα (ESSRA)* and ERRα target genes (including *UQCRSF1*, *NDUF5A*, *SDHA*, *SDHB*) after constitutive expression of PMVK in shCtrl, shFDFT1#2 and shFDFT1#3-expressing cells. Mean +/- SEM of n=3 independent biological replicates. One-way paired ANOVA test. **B.** RT-qPCR of *ERRα*, *UQCRSF1* and *NDUF5A* genes in Ctrl and PMVK-expressing cells treated every 2 days with the ERRα inhibitor XCT-790. Mean +/- SEM of n=2 independent biological replicates. **C.** RT-qPCR of *PMVK* gene in Ctrl and PMVK-expressing cells, either previously transfected by siRNA against ERRα (siERRα) (left panel) or treated every 2 days with the ERRα inhibitor XCT-790. Mean +/- SEM of n=2-3 independent biological replicates. One-way paired ANOVA test. **D.** Growth curves of Ctrl and PMVK-expressing cells treated every 2 days with the ERRα inhibitor XCT-790. Mean +/-SEM of n=3 independent biological replicates. Paired Student's T-test on last time point. **E.** Crystal violet staining at day 12 of PMVK-expressing cells treated or not with XCT-790. **F.** Quantification of SA-β-gal positive cells in Ctrl and PMVK-expressing cells treated or not with XCT-790. Mean +/-SEM of n=3 independent biological replicates. One-way paired ANOVA test. **G.** RT-qPCR of $p21^{CIP1}$ and *IL-8* genes in Ctrl and PMVK-expressing cells treated with XCT-790. Mean +/-SEM of n=

2 independent biological replicates. One-way paired ANOVA test. **H.** RT-qPCR of ERRα target genes (*Uqcrfs1*, *Nduf5a*, *Sdha*, *Sdhb*) in liver of ERRα WT and ERRα KO mice fed either by chow diet (CD) or high-fat diet (HFD). Mean +/- SEM of n=4 mice. One-way paired ANOVA test. **I.** Representative micrographs of liver slices stained with red oil of ERRα WT and ERRα KO mice fed either by chow diet (CD) or high-fat diet (HFD). Scale bar: 100µm.

### Supplemental Figure 1, Ziegler et al

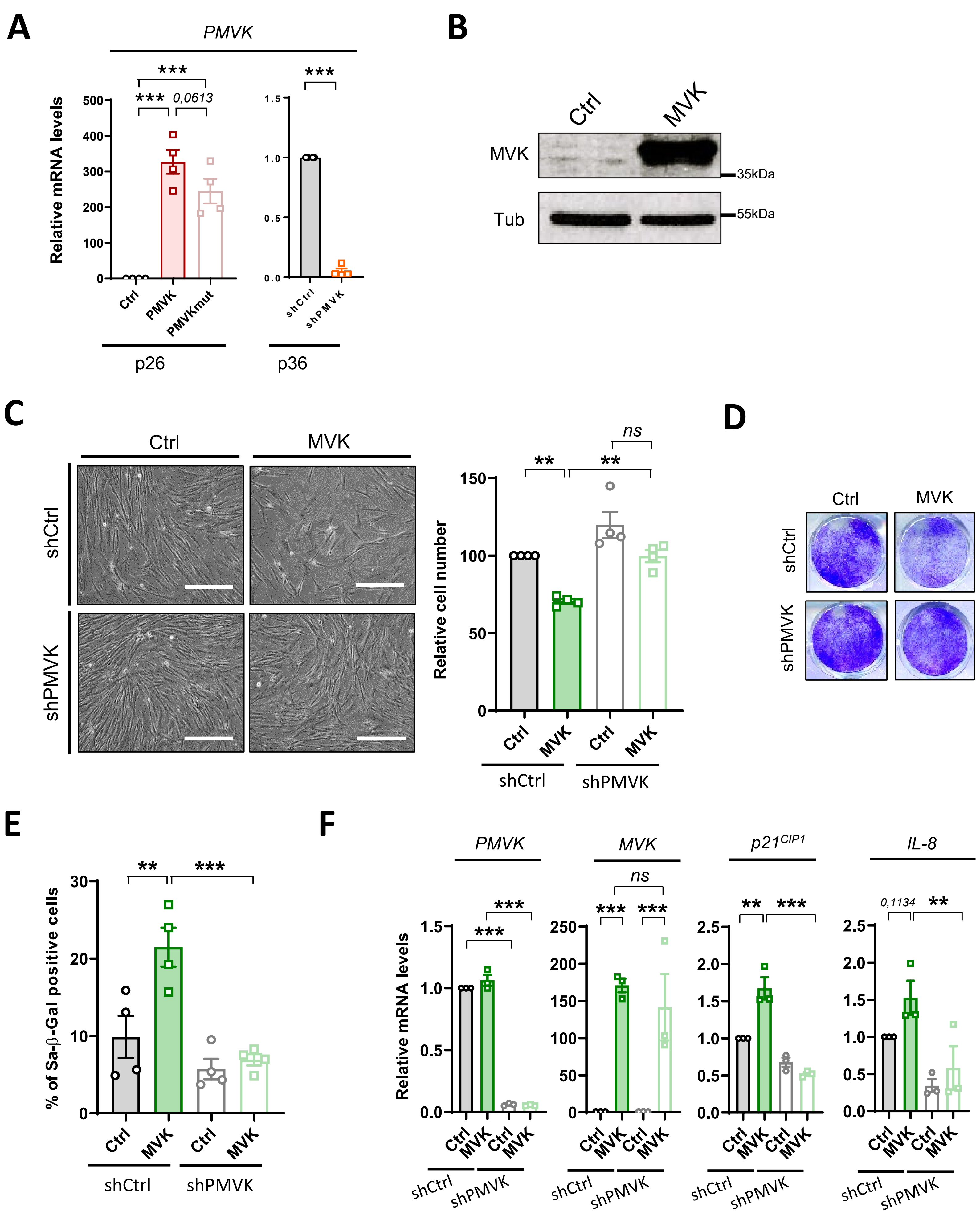

### Supplemental Figure 2, Ziegler et al

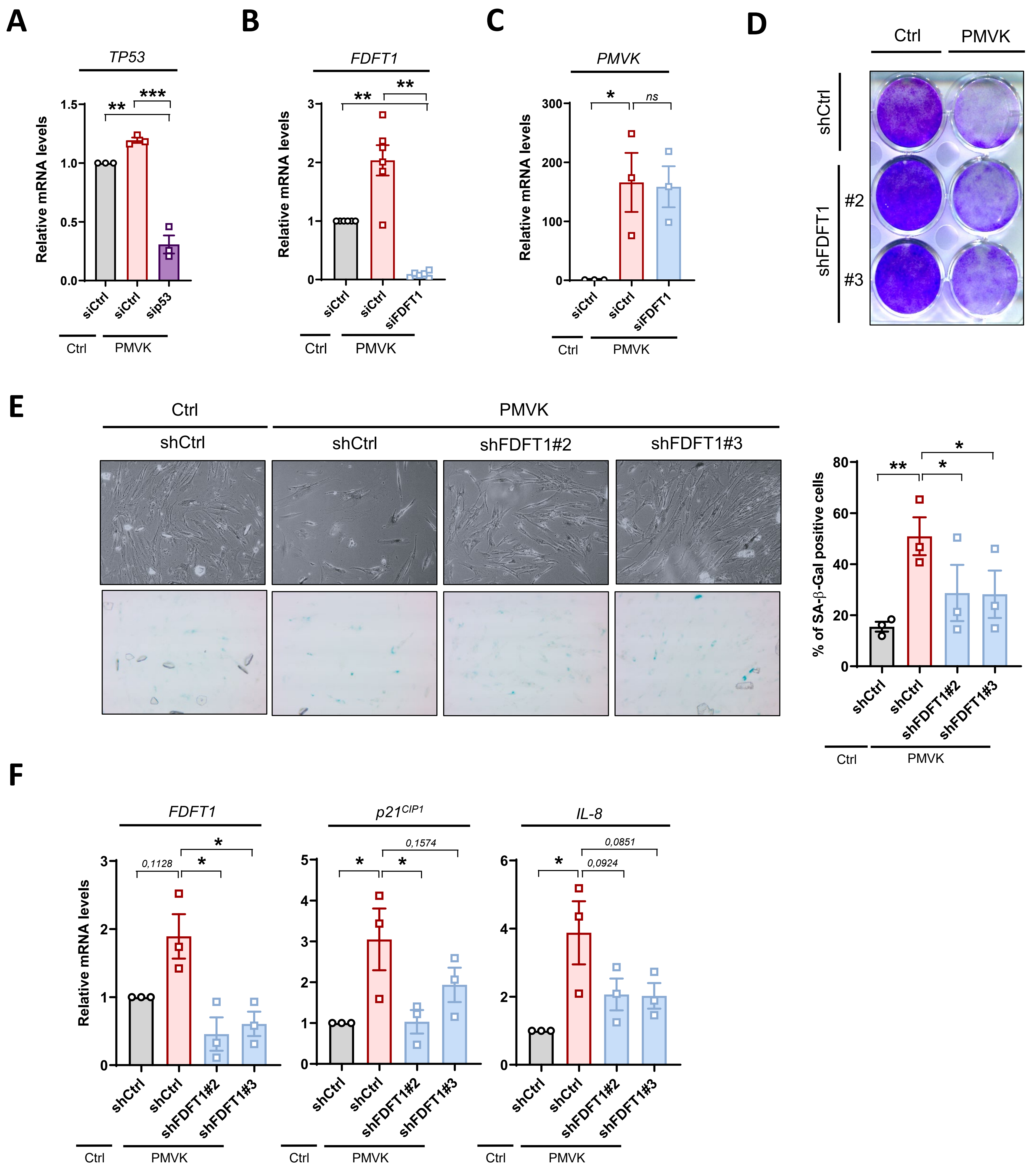

### Supplemental Figure 3, Ziegler et al

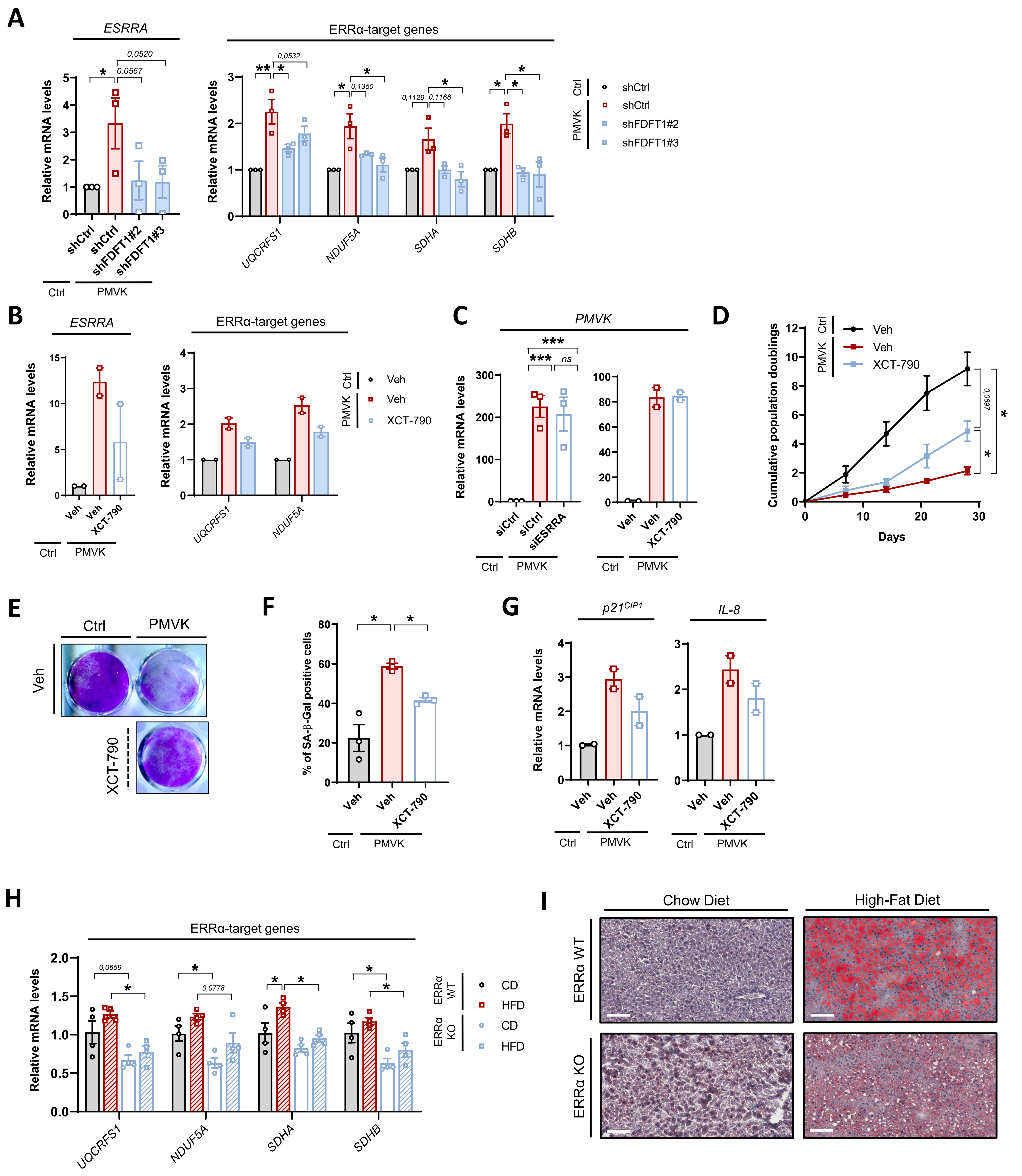

### Supplemental Table 1, Ziegler et al

| Gene | Sequence and UPL probes |
| --- | --- |
| ***Mouse*** | |
| Rplp-Forward | GCAGCAGATCCGCATGTCGCTCCG |
| Rplp-Reverse | GAGCTGGCACAGTGACCTCACACGG |
| $p21^{CIP1}$-Forward | TGAGCTGCGCCAGCTGAGGTGTG |
| $p21^{CIP1}$-Reverse | TGCCGCATGGGTTCTGACGGAC |
| Cxcl1-Forward | GGGGCGCCTATCGCCAATGA |
| Cxcl1-Reverse | AAGGCAAGCCTCGCGACCAT |
| Cxcl2-Forward | GAGCTTGAGTGTGACGCCCC |
| Cxcl2-Reverse | TCTTCCGTTGAGGGACAGCAGC |
| Gadd45a-Forward | GCTCGGAGTCAGCGCACCAT |
| Gadd45a-Reverse | GCCACATCCCGGTCGTCGTC |
| Gdf15-Forward | TGCATGCCAACCAGAGCCGA |
| Gdf15-Reverse | GACCCTGACTCAGCGACGCC |
| Uqcrfs1-Forward | ATGTGAAGCGACCCTTCCT |
| Uqcrfs1-Reverse | ATGGGAAAAACGGACAGAAG |
| Nduf5a-Forward | TGGAAGAGGTGATTCTTCAGG |
| Nduf5a-Reverse | TATTGGCCACTTCCACTGGT |
| Sdha-Forward | CAGTTCCACCCCACAGGTA |
| Sdha-Reverse | TCTCCACGACACCCTTCTGT |
| Sdhb-Forward | CTGGTGGAACGGAGACAAGT |
| Sdhb-Reverse | GCGTTCCTCTGTGAAGTCGT |
| ***Human*** | |
| ACTB-Forward | ATTGGCAATGAGCGGTTC |
| ACTB-Reverse | GGATGCCACAGGACTCCAT |
| $p21^{CIP1}$-Forward | TCACTGTCTTGTACCCTTGTGC |
| $p21^{CIP1}$-Reverse | GGCGTTTGGAGTGGTAGAAAT |
| MVK-Forward | GCCTTTCTTTACTTATACCTGTCCA |
| MVK-Reverse | CCGACCACACTACGATATCCA |
| PMVK-Forward | TTTTGCAGGAAGATTGTGGA |
| PMVK-Reverse | CTCCGTGTGTCACTCACCA |
| IL8-Forward | AGACAGCAGAGCACACAAGC |
| IL8-Reverse | ATGGTTCCTTCCGGTGGT |
| CXCL1-Forward | TCCTGCATCCCCCATAGTTA |
| CXCL1-Reverse | CTTCAGGAACAGCCACCAGT |
| TP53-Forward | AGGCCTTGGAACTCAAGGAT |
| TP53-Reverse | CCCTTTTTGGACTTCAGGTG |
| GADD45A-Forward | AGAGCAGAAGACCGAAAGGA |
| GADD45A-Reverse | TGACTCAGGGCTTTGCTGA |
| GDF15-Forward | CCGGATACTCACGCCAGA |
| GDF15-Reverse | AGAGATACGCAGGTGCAGGT |
| FDFT1-Forward | AGTTTCGCAGCTGTTATCCAG |
| FDFT1-Reverse | GATAAAATATGCACACTGCGTTG |
| ESSRA-Forward | GGCGGCAGAAGTACAAGC |
| ESSRA-Reverse | ATTCACTGGGGCTGCTGT |
| UQCRFS1-Forward | AGCCTGTGTTGGACCTGAAG |
| UQCRFS1-Reverse | TGGGAATAACAAACAGAAGCAG |
| NDUF5A-Forward | GGTGTGCTGAAGAAGACCACT |
| NDUF5A-Reverse | TTGTGTACAATATTCTTAGCCTCTCG |
| SDHA-Forward | TCCACTACATGACGGAGCAG |
| SDHA-Reverse | CCATCTTCAGTTCTGCTAAACG |
| SDHB-Forward | GGGGCCTGCAGTTCTTATG |
| SDHB-Reverse | AGGCGCTCCTCTGTGAAGT |

### Supplemental Table 2, Ziegler et al

| Protein | Reference | Use | Dilution |
|---|---|---|---|
| MVK | sc-27585 (Santa-Cruz) | WB Figure 1 | 1/500 |
| PMVK | sc-390775 (Santa-Cruz) | WB Figure 1 | 1/500 |
| Tub | T6199 (Sigma-Aldrich) | WB Figure 1 | 1/5000 |
| GammaH2AX | 2577S (Cell Signaling) | IF Figure 2 | 1/300 |
| ERRα | Ab76228 (Abcam) | WB Figure 4 | 1/1000 |
| $p21^{CIP1}$ | Sc-817 (SantaCruz) | WB Figure 4 | 1/1000 |
| Tub | CLT9002 (Cedarlane) | WB Figure 4 | 1/2000 |
